# Time-dependent enhancement in ventral tegmental area dopamine neuron activity drives pain-facilitated fentanyl intake in males

**DOI:** 10.1101/2022.08.19.504549

**Authors:** Jessica A. Higginbotham, Julian G. Abt, Rachel H. Tiech, Jose A Morón

**Affiliations:** Department of Anesthesiology, Washington University in St. Louis, St. Louis, MO, USA; Pain Center, Washington University in St. Louis, St. Louis, MO, USA; School of Medicine, Washington University in St. Louis, St. Louis, MO, USA; Department of Neuroscience, Washington University in St. Louis, St. Louis, MO, USA; Department of Psychiatry, Washington University in St. Louis, St. Louis, MO, USA

## Abstract

Pain affects over 50% of US adults. Opioids are potent analgesics used to treat pain symptoms but are highly prone to abuse – creating a major dilemma for public health. Evidence suggests that the proclivity for opioid abuse under pain conditions varies between sexes. However, the neural mechanisms underlying sex-specific effects of pain on opioid use are largely unclear. Here, we recapitulate clinical findings and demonstrate that pain increases self-administration of the widely abused opioid, fentanyl, selectively in male rats. These behavioral effects develop over time and are paralleled by sex- and pain-specific effects on fentanyl-evoked ventral tegmental area (VTA) dopamine (DA) neuron activity, a critical mediator of motivation and reward. Using *in vivo* fiber photometry, we show that tonic VTA DA neuron activity is attenuated in males with pain. In contrast, phasic VTA DA neuron responses to self-administered fentanyl increase in magnitude at later timepoints and correspond with increases in fentanyl intake. The protracted increase in fentanyl-evoked VTA DA activity is necessary for pain to enhance fentanyl self-administration in males because chemogenetic inhibition of VTA DA neurons normalized fentanyl intake and associated fentanyl-evoked VTA DA neuron responses. These findings reveal time-dependent and sex-specific pain-induced adaptations to VTA DA neuron function that underlie maladaptive patterns of opioid use.

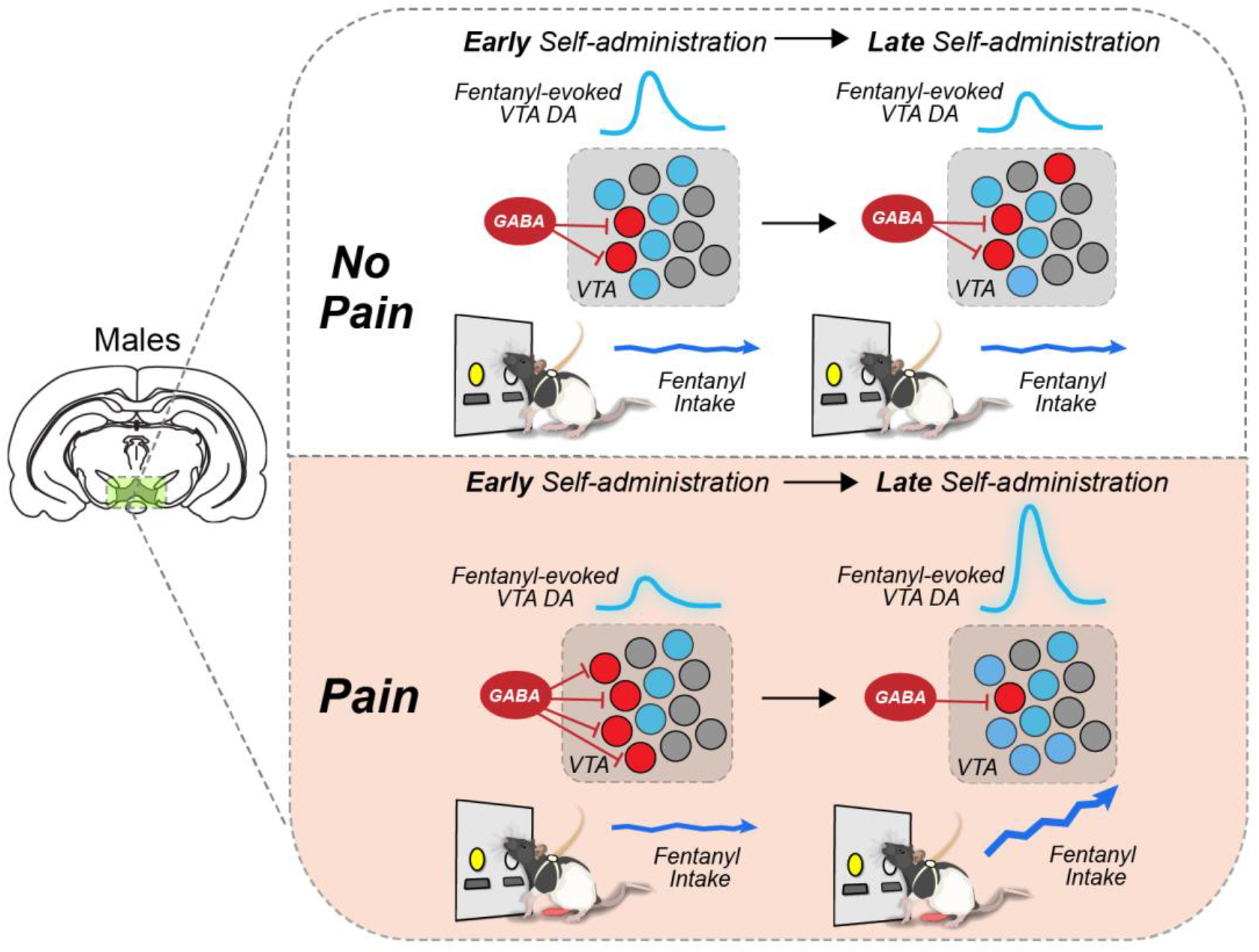

## INTRODUCTION

The intersection between pain and opioid abuse presents a major dilemma for public health. Over half of US adults report having pain^1^, which is a primary source of motivation for opioid misuse^2^. Opioid pain medications are potent analgesics commonly used to treat pain symptoms but are highly prone to abuse. Consequently, efforts to mitigate opioid use through prescribing practices have been met with alarming increases in opioid overdose deaths^3^. Interestingly, clinical evidence suggests that the effects of pain on opioid misuse liability are gender/sex specific^4^. Although women are more sensitive to pain^5^, men are more likely to escalate opioid doses^6^, misuse prescriptions ^6,7^, and meet diagnostic criteria for opioid use disorder^8^. Despite this, our understanding of sex differences underlying the effects of pain on opioid use remains limited. As such, revealing neural mechanisms underlying sex-specific effects of pain on opioid abuse is of timely importance from a pain treatment perspective.

The mesolimbic dopamine (DA) pathway plays a prominent role in reward processing and motivated behaviors through phasic ventral tegmental area (VTA) DA release into the nucleus accumbens (NAc)^9–11^. Opioids trigger large increases in extracellular DA in the NAc to produce their reinforcing effects^12–14^. Importantly, pain negatively impacts the ability of opioids to trigger mesolimbic DA release, which may contribute to opioid misuse liability. Human imaging studies demonstrate that pain reduces mesolimbic DA pathway connectivity and DA receptor binding in the NAc^15–18^. Similarly, preclinical models of inflammatory or neuropathic pain exhibit reduced VTA DA neuron excitability and opioid evoked DA release in the NAc^19,20^. We previously found that heroin administration in rats with inflammatory pain at low or high doses produces deficits or enhancements in extracellular NAc DA release, respectively^20^. Moreover, the effects of pain on NAc DA release are associated with similar effects on self-administration such that, rats with pain self-administer less heroin at low doses and more heroin at higher doses^20^, but these effects were only evaluated in males. While this evidence indicates that pain perturbs mesolimbic DA function and alters opioid self-administration, it is unknown whether pain disrupts DA activity and opioid self-administration in a sex-specific manner. Moreover, the extent to which opioid-naïve rats with pain develop maladaptive patterns of opioid use and alterations in opioid-evoked VTA DA activity remains unclear.

While current evidence supports that pain-induced alterations in mesolimbic DA function lead to altered motivational states^20–23^, no studies to date have considered whether this phenomenon is sufficient to drive aberrant opioid use in a sex-specific manner. Women experience pain with higher frequency and intensity^24,25^, but are less sensitive to the analgesic effects of opioids^26–28^. Paradoxically, men are more likely to engage in prescription opioid misuse and dose escalation^6,8,29–31^ and are twice as likely die from overdose^32^. Sex differences are also observed in within the mesolimbic DA system. DA neurons are larger and more abundant in the VTA of females compared to males^33,34^. Moreover, females have higher rates of DA release and reuptake in the NAc, which can be sensitive to circulating ovarian hormones^35–37^. Therefore, sex differences in mesolimbic DA system function under conditions of pain may influence the trajectory and expression of maladaptive opioid use.

Using wireless *in vivo* fiber photometry combined with intravenous fentanyl self-administration, we report that male rats with persistent inflammatory pain exhibit time-dependent increases in fentanyl consumption and accompanying fentanyl-evoked VTA DA neuron calcium transients. These effects of pain are specific to males and driven by pain-facilitated motivation for fentanyl reward at higher doses rather than sex differences in inflammation, nociception, or antinociception. Finally, we demonstrate that the protracted effects of pain on increased fentanyl consumption in males necessitate corresponding increases in fentanyl-evoked VTA DA neuron response activity by chemogenetically inhibiting VTA DA neuron activity during late stages of self-administration. Together, these results indicate that pain induces sex-specific neuroadaptations within VTA DA neuron function that contribute to differences in opioid reward encoding within the mesolimbic DA pathway underlying increased opioid intake. Our findings provide novel insight as to how pain alters opioid reward signaling in a sex-specific and time-dependent manner and reveal novel mechanisms underlying opioid misuse liability.

## RESULTS

### Pain increases self-administration of fentanyl in males in a time-dependent manner

To examine whether pain alters opioid intake in a sex-specific manner, we used complete Freund’s adjuvant (CFA) to mimic persistent inflammatory pain in male and female rats and an instrumental model of drug self-administration. After intravenous catheterization and intraplantar injections with CFA (males, 150 nL; females 120 nL) or SAL (males, 150 nL; females, 120 nL) in the right hind paw, rats were trained on a fixed-ratio 1 (FR1) schedule of reinforcement to self-administer fentanyl during 5 daily 2-hr sessions/week. During the first week, rats were trained with fentanyl infusion doses of 5 μg/kg (i.v.) to facilitate acquisition, and doses of 2 μg/kg (i.v.) were used during successive weeks (**Fig. 1A**). Despite the lack of training history prior to initiating fentanyl-self administration, rats exhibited bias toward the correct (fentanyl-paired) lever from the first day of training (**Fig. 1B-C and Extended Data Fig. 1A**). As such, pain and sex did not influence the ability of rats to acquire fentanyl self-administration or discriminate between levers.

**FIGURE 1:**
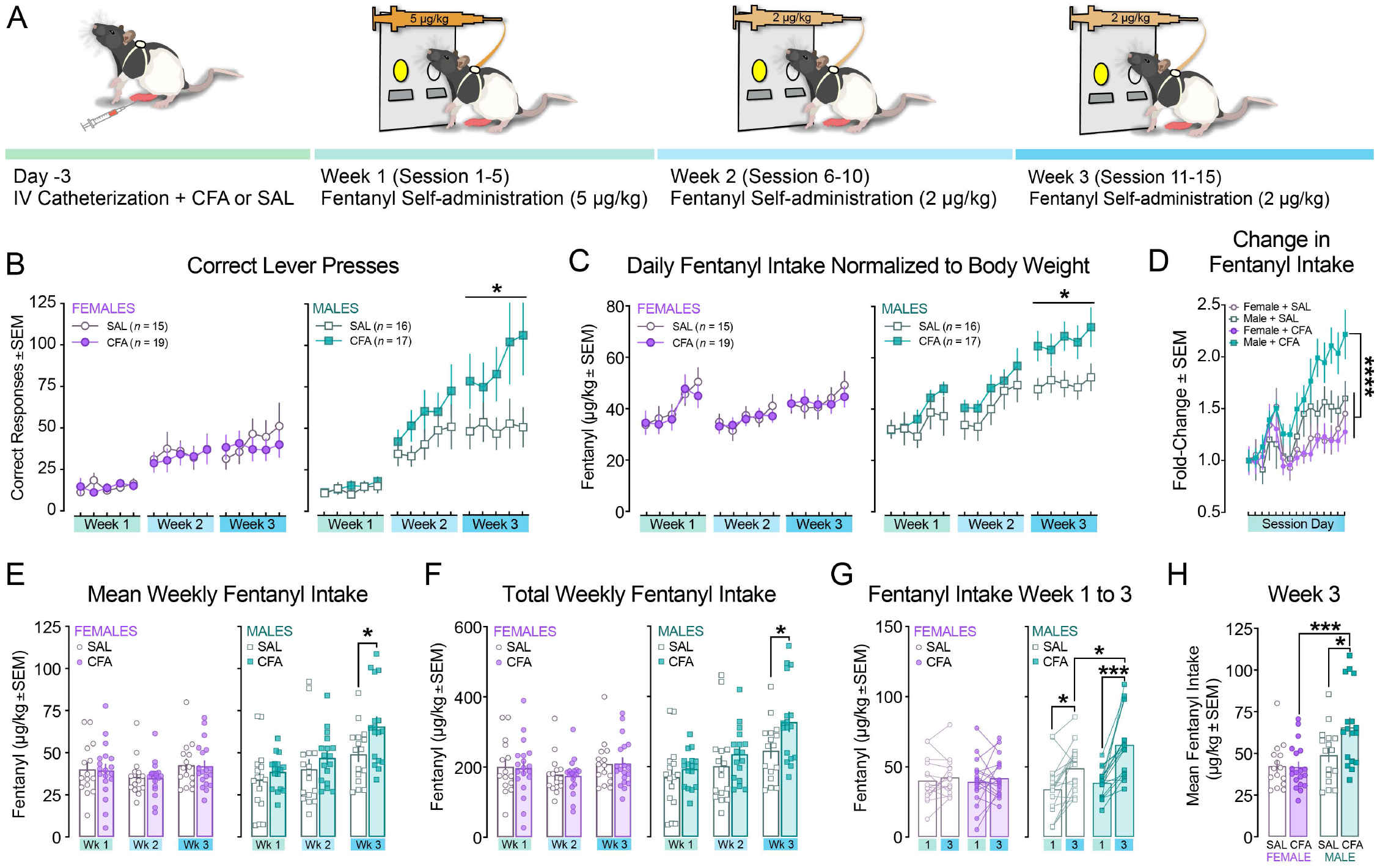
Pain increases fentanyl consumption selectively in males. **A,** Schematic representation of behavioral methodology. **B,** In females (*left*), pain does not alter fentanyl responses in females (RM 2way ANOVA, time: F_(4, 128)_ = 2.080, P=0.087; time x pain: F_(4, 128)_ = 2.418, P=0.052; n=15 (SAL, open symbols) or 19 (CFA, closed symbols). In males (*right*), pain increases responding for fentanyl during week three (RM 2way ANOVA, time: F_(4, 120)_ = 3.097, P=0.018; time x pain: F_(4, 120)_ = 2.621, *P=0.038; n=16 (SAL, open symbols) or 17 (CFA, closed symbols). **C,** Pain does not affect fentanyl intake in females (*left*), but increases fentanyl intake (normalized to body weight) in males during week 3 (*right;* RM 2way ANOVA, pain: F_(1,31)_=5.490, *P=0.026). **D,** Males with pain increase fentanyl intake (intake/first session intake, μg/kg) compared to males with SAL or females with CFA (RM one-way ANOVA: F_(3,42)_ = 30.79, P<0.0001; Holm-Sidak’s post hoc, ****P<0.0001). **E,** Mean weekly fentanyl intake (μg/kg) is similar across weeks in females with or without pain (*left*). Pain increases mean fentanyl consumption in males during week 3 (*right;* RM 2way ANOVA, time: F_(2,63)_=33.66, P<0.0001; Sidak’s post hoc, *P=0.043). **F,** Total weekly fentanyl intake (μg/kg) is similar between females with pain or no pain (*left*) and higher in males with pain compared to no pain during week three (*right;* RM 2way ANOVA, time x pain: F_(2,62_)=2.997, P=0.057; time: F_(2,62)_=33.25; Sidak’s post hoc, *P=0.033). **G,** Females do not increase fentanyl intake over time (*left*; 2way ANOVA, time: F_(1,32)_=1.032, P=0.317). Males increase fentanyl intake between weeks 1 and 3 (*right*; 2way ANOVA, time: F_(1,31)_=24.34, P<0.0001, Sidak’s Post Hoc, **P=0.005, ****P<0.0001) and males in pain take more fentanyl during week 3 than males without pain (Sidak’s post hoc, *P=0.020). **H,** Males with pain have higher fentanyl consumption during the third week of self-administration than males without pain or females with pain (2way ANOVA: Sex x pain: F_(1,63)_=4.092, P=0.047; Sidak’s post hoc, *P=0.015, ***P=0.0002).

Fentanyl consumption during weeks 1 and 2 were unaffected by the presence of pain (CFA), but during week 3, males with pain significantly increased their responding for fentanyl and fentanyl intake (normalized to body weight) relative to males without pain (SAL), while females maintained similar levels of responding across weeks regardless of treatment (**Fig. 1B-C**). The ratio of intake on a given day relative to the first session showed that males with CFA increased their intake of fentanyl (μg/kg) relative to males with SAL or either female group (**Fig. 1D**). Mean and cumulative weekly fentanyl consumption was stable over time in females with CFA and SAL (**Fig. 1E-G**). In contrast, males showed time-dependent increases in weekly fentanyl self-administration, which was exacerbated by pain during week 3 (**Fig 1E-G**). Males with pain took more fentanyl during the last week of self-administration than females with pain or males without pain (**Fig. 1H**) indicating that the facilitative effects of pain on fentanyl consumption were both sex-specific and time-dependent. This evidence supports the idea that pain may yield males more vulnerable to maladaptive patterns of opioid use under conditions of long-term opioid exposure.

### CFA-induced inflammation, nociception, and opioid-evoked analgesia are similar between sexes and groups throughout fentanyl self-administration

To begin to dissect potential mechanisms by which pain selectively increases fentanyl consumption in males, we assessed whether pain had sex-specific effects on CFA-induced inflammation, nociception, and antinociception (**Fig. 2A**). To determine whether pain produced by CFA manifested differently between sexes or changed over the three weeks of fentanyl self-administration, we measured relative paw thickness and mechanical sensitivity in a subset of rats over the course of the experimental timeline. CFA-injected paws were about twice the size of the contralateral non-injected paw in both sexes which was maintained throughout training (**Fig. 2B**). Nociceptive responses were evaluated using relative paw withdrawal thresholds (PWT) in response to mechanical stimulation. Relative PWT obtained from injected paws were similar between sexes within treatment groups and were stable across time (**Fig. 2C**), negating the possible development of fentanyl-induced hyperalgesia during training. These results demonstrate that CFA produced comparable levels of inflammation and nociceptive responses between sexes and are consistent with previous reports indicating the stability of this approach^38^. Tolerance, a progressive reduction in drug efficacy, is known to facilitate opioid use^39–41^. We assessed this possibility at least two hours after the fentanyl self-administration session by comparing baseline PWT to PWT following a single fentanyl infusion (2 μg/kg, i.v.). This dose was sufficient to produce analgesia (indicated by increased PWT) and the magnitude of the effect was overall, greater in pain conditions, but showed minimal variability over time between sexes (**Fig. 2D**). Together, these findings demonstrate that males with pain do not increase fentanyl intake due to sex-specific or time-dependent effects on CFA-induced inflammation, nociception, or antinociception.

**FIGURE 2:**
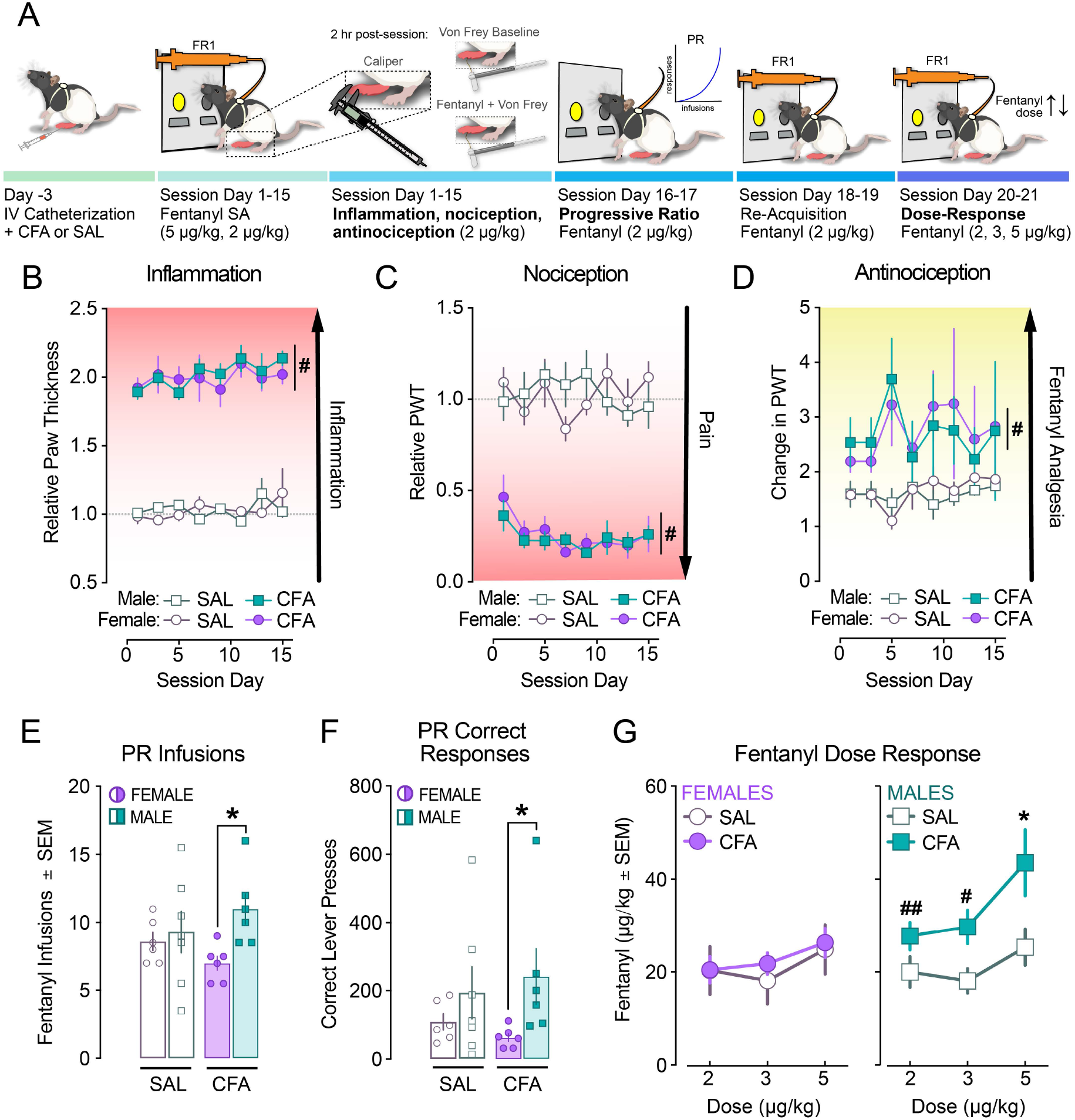
Males and females exhibit similar inflammation, nociception, and antinociception while pain has sex-specific effects on motivation and reinforcement. **A,** Schematic representation of behavioral methodology. **B,** Caliper measurements of paw thickness (mm) after CFA produces similar levels of relative inflammation (injected paw/ non-injected paw) in males and females throughout self-administration (RM 3way ANOVA, sex x pain: F_(1, 24)_=0.3138, P=0.581; pain: F_(1, 24)_=950.6, #P<0.0001). **C,** Pain sensitivity measured as paw withdrawal threshold (PWT) of injected paw relative to non-injected paw is similar between males and females (RM 3way ANOVA, sex x pain: F_(1,167)_=1.659, P=0.1995; pain: F_(1,167)_=122.4, #P<0.0001). **D,** Analgesia produced by a single fentanyl infusion (2 μg/kg, i.v.) was similar between males and females within treatment groups (RM 3way ANOVA, sex x pain: F_(1,24)_=0.0019, P=0.966; pain: F_(1,24)_=21.58, #P<0.0001). **E,** Pain increases motivation for fentanyl, indicated by higher break points, in males relative to females (2way ANOVA, sex: F_(1,21)_=4.497, P=0.046; Sidak’s Post hoc, *P=0.041). **F,** Males have higher motivation for fentanyl, indicated by a higher number of correct responses during the progressive ratio (PR) tests (2way ANOVA, sex: F_(1,21)_=4.760, #P=0.041). **G,** In females, pain does not alter fentanyl dose-response (left; 2way ANOVA, dose: F_(2,16)_=4.254, P=0.033; pain: F_(1,8)_=0.1045, P=0.755), but males with pain take more fentanyl at the highest dose (5 μg/kg) than any other dose (2-3 μg/kg) and more than males without pain at the same dose (right; 2way ANOVA, dose: F_(2,16)_=8.452, P=0.003; between-subjects, Sidak’s Post hoc, *P=0.036; within-subjects, Sidak’s Post hoc, #P<0.05, ##P<0.01).

### Males with inflammatory pain are motivated by higher fentanyl doses

We assessed whether the effects of pain on fentanyl self-administration were due to sex differences in the motivation for fentanyl, fentanyl-paired cues, or reinforcing properties of fentanyl. To do this, we examined three different components of instrumental drug use: extinction/cue-induced reinstatement of drug-seeking behavior, motivated behavior (progressive ratio), and reinforcing properties of fentanyl (dose-response). To examine if pain influenced persistent drug-seeking behavior in a sex-specific manner, a subset of rats (n=12-14/group) underwent 7 days of extinction training (no fentanyl or fentanyl-paired cues) followed by a test of cue-induced reinstatement (fentanyl-paired cues without fentanyl) after completing three weeks of fentanyl self-administration (**Extended Data Fig. 1B-C**). Persistent drug-seeking behavior during extinction was higher in males with pain than no pain, but comparable to that of both female groups (**Extended Data Fig. 1B**). In contrast, cue-induced reinstatement of drug-seeking behavior was unaffected by pain, while the magnitude of reinstatement (non-reinforced lever presses) was higher in males compared to females (**Extended Data Fig. 1C**). Although unaffected by pain, the sex differences in reinstatement suggest that drug-paired cues exude higher motivational salience over males than females.

To evaluate motivated behavior for fentanyl self-administration, progressive ratio (PR) tests were administered^42^ to a separate cohort on two consecutive days during which each response requirement for a fentanyl infusion (2 μg/kg, i.v.) increased exponentially during the successive trial throughout the 2-hr session (**Fig. 2A**). Pain increased the mean number of fentanyl infusions obtained (**Fig. 2E**) and the number of responses in males compared to females (**Fig. 2F**). However, because males responded similarly under pain or no pain conditions, these findings suggest that males are resistant to motivational deficits produced by pain. To examine whether this could be associated with sex-dependent effects of pain on the reinforcing properties of fentanyl, we conducted two inter-session dose-response tests with counter-balanced ascending or descending doses of fentanyl (2,3,5 μg/kg/infusion, i.v.). In females, the mean intake at each dose trended toward preference of the highest dose, but CFA did not affect intake at any dose (**Fig. 2G**). In contrast, males with CFA increased fentanyl intake at the highest available dose (5 μg/kg, i.v.) relative to other available doses (2-3 μg/kg, i.v.) and compared to males with SAL (**Fig. 2H)**. The high fentanyl dose preference of males with pain aligns with our previous reports showing high heroin dose preference in rats 3 days after hind paw injections of CFA (females not evaluated)^20^.

### VTA DA cell activity is impacted by pain in a sex-specific and time-dependent manner

Opioids stimulate VTA DA release through mu-opioid receptor-dependent disinhibition. Previous work in our lab found increased inhibitory drive onto VTA DA neurons three days after hind paw CFA injections^20,23^. This effect is also associated with altered ability of opioids to stimulate NAc DA release which paralells dose-dependent effects of CFA on heroin motivation^20^. Thus, we hypothesized pain increases fentanyl use in males due to sex differences in the ability of fentanyl to stimulate VTA DA cell responses. To test this, we combined intravenous fentanyl self-administration with wireless *in vivo* fiber photometry to examine in real time how pain effects instrumental fentanyl-evoked VTA DA activity. Exploiting on this technique, we examined how this response was impacted at multiple timepoints throughout the three weeks of self-administration to assess potential adaptations to fentanyl-evoked responses associated with enhanced drug-taking behavior.

To selectively measure VTA DA cell calcium transient activity, we injected a Cre-dependent calcium sensor (pGP-AAV-syn-FLEX-jGCaMP7c-WPRE) and implanted an optic fiber in the VTA of male and female rats expressing Cre-recombinase in tyrosine hydroxylase (TH) positive neurons (TH-Cre+; **Fig. 3A-B, Extended Data Fig. 2**). As before, rats were implanted with IV catheters and injected in the right hind-paw with CFA or SAL before fentanyl self-administration training. Intravenous fentanyl self-administration and detection of fentanyl-evoked calcium transients from VTA DA neurons (TH-Cre+) were achieved in freely behaving rats using a wireless photometry device^43,44^. GCaMP fluorescence VTA-infected cells were recorded at from each rat for the duration of the 2-hr session at least once per week for each of the three weeks of training. On the first day of recording, the device was secured to the rats’ indwelling optic fiber and the LED power (<36 mW) was adjusted to achieve optimal output (30-40k a.u.). The settings determined during the first recording session were used during each subsequent recording session to produce stable baseline fluorescent signals for up to three weeks (i.e., duration of experiment; see **Extended Data Fig. 3**). Tonic VTA DA calcium dynamics were sampled at 10 Hz throughout the 2-hr session while phasic events were aligned to fentanyl lever responses and measured for 30 s (−10 to +20 s relative to response).

**FIGURE 3:**
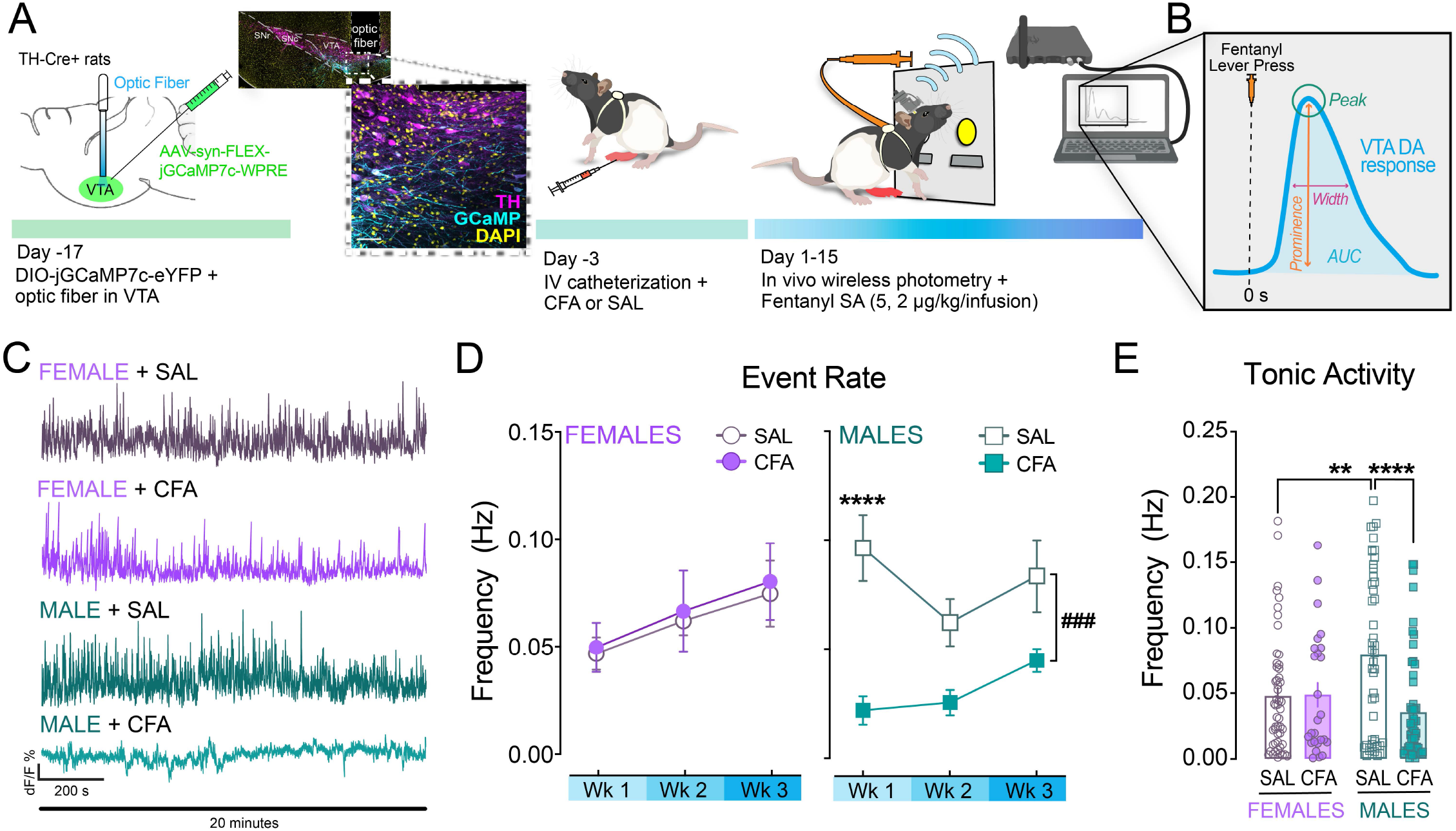
Wireless *in vivo* photometry detects sex-specific effects of pain on tonic VTA DA calcium transient activity. **A,** Schematic of behavioral methodology with wireless *in vivo* photometry and representative image of fiber placement and GCaMP expression co-localized with TH in the VTA of TH-Cre+ rats (scale bar = 50 μm). **B,** Schematic depicting event determinants. For tonic activity, events were measured as peaks (green circle) with prominence (rise above baseline) of at least 1.2x dF/F and at least 150 ms in width at half-prominence. Phasic VTA DA cell responses were aligned to lever presses reinforced with fentanyl (orange syringe, T=0s) and peaks above threshold were at least 1.2x dF/F in prominence and at least 500 ms in width at half-prominence. Area under the curve (AUC) was measured at −4 to +8 s relative to lever press and binned in 2-s intervals. **C,** Representative traces depicting tonic activity of VTA TH+ cells during self-administration **D,** Frequency of events detected throughout the recording sessions were similar over time in females (*left*) with pain (CFA, filled circles) or no pain (SAL, open circles). In males (*right*), pain (CFA, filled squares) reduced the frequency of events across the three weeks of self-administration compared to males without pain (SAL, open squares) (RM 2way ANOVA; time x pain: F_(2,24)_ = 3.710, P=0.039; pain: F_(1,15)_ = 16.59, ###P=0.0010; time: F_(2,24)_=4.323, P=0.025; Sidak’s post hoc, ****P<0.0001) **E,** In the absence of pain (SAL), males exhibit increases calcium transient event frequency, while pain (CFA) reduces frequency in males but not females (2way ANOVA, sex x pain: F_(1,178)_=8.534, P=0.0039; pain: F_(1,178)_=7.807, P=0.0058; Sidak’s post hoc, *P=0.011, ****P<0.0001).

We examined whether tonic VTA DA calcium transient activity was different between sexes or impacted by pain by measuring changes in the local fluorescent maxima with typical GCaMP kinetics to define spontaneous transient events throughout the 2-hr self-administration session^23^ (**Fig. 3B).** Differences in the frequency of VTA DA events were dependent on pain and varied between sexes (**Fig. 3C-E**). VTA DA event frequency was similar in females independent of pain, but increased over time overall (**Fig. 3D**). In males, pain reduced event rates compared to no pain, despite increasing in frequency over time (**Fig. 3D**). Overall, males without pain had higher frequency VTA DA neuron activity than females or males with pain (**Fig. 3E)**. These results indicate that pain reduces the overall frequency of calcium transients in VTA DA neurons in a sex-specific manner and are consistent with the idea that pain reduces tonic activity of VTA DA neurons^21^.

To examine whether pain had time-dependent and sex-specific effects on VTA DA cell calcium transients evoked by self-administered fentanyl, photometry recording signals were time-locked to each fentanyl-reinforced lever press (trial) (**Fig. 4A, E)**. In females, fentanyl evoked responses from VTA DA cells that were similar in magnitude and stable over time (**Fig. 4A-D).** In females injected with CFA, AUC were similar to SAL during the first week of self-administration but was maintained over time (**Fig. 4B-C)** despite subtle differences in the shape of the response (**Fig. 4B and Extended Data Fig. 4A-C).** The change in overall shape resulted in reduced total AUC of fentanyl-evoked calcium transients that appeared over time in females without pain relative to pain (**Fig. 4C**). For each trial, the fentanyl-evoked peak probability was calculated based on the ability of self-administered fentanyl infusion to trigger a significant increase VTA DA transient signal^23^. In females, fentanyl-evoked VTA DA peak probabilities were similar throughout the three weeks of self-administration (**Fig. 4D).** Together, these results indicate that pain has minimal effects on fentanyl-evoked VTA DA activity in females.

**FIGURE 4:**
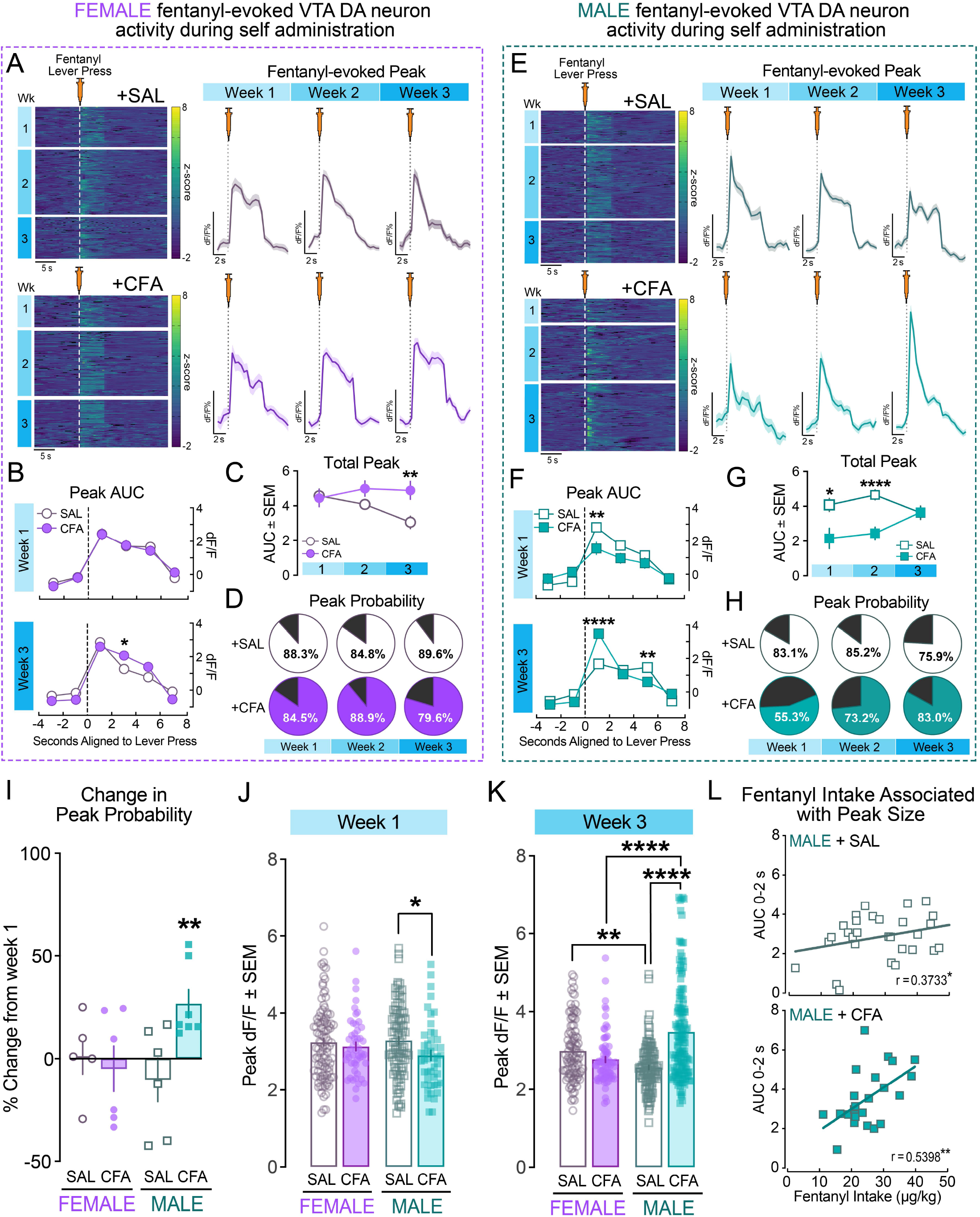
Phasic fentanyl-evoked VTA DA cell calcium transient activity is time-dependently modified in males with pain and associated with increased fentanyl consumption. **A,** Heatmaps showing weekly VTA DA cell response (z-score) from aligned to lever presses (dotted line) resulting in a fentanyl infusion and averaged photometry traces showing responses to fentanyl (syringe) in females without pain (SAL; n=5 rats, 4-28 trials/recording session, 1-2 recording sessions per week) or with pain (CFA; n=6 rats, 4-24 trials/recording session, 1-2 recording sessions per week). **B,** In females, pain does not change area under the curve (AUC) of phasic VTA DA cell responses during week 1 and week 3 responses from females with pain (CFA, filled circles) are slightly larger 2-4 s after the lever press than females without pain (SAL, open circles), but otherwise unaffected (2way ANOVA, time x treatment: F_(5,1344)_=4.749, P=0.0003: time, F_(5,1344)_=105.2, P<0.0001; Sidak’s post hoc, *P=0.038). **C,** Total AUC across weeks is higher in females with pain (CFA; filled circles) than no pain (SAL; open circles) during week 3 (2way ANOVA, treatment: F_(1,594)_=6.593, P=0.011; Sidak’s Post hoc, **P=0.007).**D,** The percentage of obtained rewards that evoked a response was not different between female groups across time. **E,** Heatmaps by week of GCaMP z-scores from each trial where a lever press (dotted line) resulted in a fentanyl infusion and mean dF/F signals from each week of self-administration aligned to reward (syringe) in males without pain (SAL; n=6 rats, 5-32 trials/recording session,1-2 sessions per week) or with pain (CFA; n=7 rats, 5-36 trials/recording session, 1-2 sessions per week). **F,** Pain (CFA; filled squares) reduces AUC of phasic VTA DA cell responses 0-2 s after the lever press compared to males without pain (SAL; open squares) during week 1 of self-administration (2way ANOVA, time x pain: F_(2,535)_=13.79, P<0.0001; pain: F_(1,535)_=18.75, P<0.0001; Sidak’s Post hoc, ##P<0.01 vs. Week 3), but during week three, decay effects of CFA on fentanyl evoked AUC are still reduced but initial responses were higher than SAL (2way ANOVA, time x pain: F_(5,1554)_=15.27, P<0.0001; time: F_(5,1554)_=107.9, P<0.0001; Sidak’s post hoc, **P=0.0023, ****P<0.0001). **G,** Total AUC across weeks is higher in males without pain (SAL; open squares) than no pain (CFA; filled squares) during weeks 1 and 2, but are similar during week 3 (2way RM ANOVA; time x pain: F_(2,698)_=6.294; pain: F_(1,698)_=18.81; time: F_(2,698)_=0.833; Sidak’s post hoc, *P<0.05, ****P<0.0001). **H,** The percentage of obtained rewards that evoked a response was markedly reduced in males with pain (filled pie chart) during week 1 of self-administration, but similar to males without pain (open pie chart) by week 3. **I,** Pain selectively facilitates the ability of fentanyl to evoke responses in males by week 3 of self-administration relative to week 1 (one-sample t-test, **P=0.007). **J,** Fentanyl-evoked peak dF/F is reduced in males with pain (CFA; filled cyan bar) relative to no pain (SAL; open cyan bar) during week 1 (2way ANOVA; pain: F_(1,290)_=4.734, P=0.0304; Sidak’s post hoc, *P<0.05). **K,** Peak dF/F during week 3 is higher in control females than males while males with pain (CFA) exhibit higher amplitude peaks than males without pain (SAL) or females with pain (2way ANOVA; sex x pain interaction: F_(1,438)_=41.13, P<0.0001; pain: F_(1,438)_=15.58, P<0.0001; sex: F_(1,438)_=2.558, P=0.110; Sidak’s post hoc, **P<0.01, ****P<0.0001). **L,** Mean peak size (AUC 0-2 s) is positively correlated with session fentanyl intake in males without pain (SAL, top) or males with pain (CFA, bottom) (Pearson’s correlations, *P<0.05, **P<0.01).

In contrast, CFA had robust effects on fentanyl-evoked VTA DA activity in males (**Fig. 4E-H)**. The AUC from males injected with CFA was significantly attenuated during the first (**Fig. 4F-G)** and second **(Extended Data Fig. F)** weeks of fentanyl self-administration, but increased over time, reaching an overall, similar size of SAL males by week 3 (**Fig. 4G**). The effects on AUC masked dramatic time-dependent effects of CFA on the shape of responses evoked by fentanyl, such that calcium dynamics detected during week one from CFA males were low in amplitude and longer in duration and evolved into responses short in duration and high in amplitude (**Extended Data Fig. 4D-E).** In males, pain had time-dependent and bidirectional effects on fentanyl-evoked peak probability rates such that, injection with SAL decreased the probability of fentanyl-evoked transient events each week and CFA increased these rates over time (**Fig. 4H**).

Analysis of the change in peak probability from week 1 to week 3 revealed that only males with pain (CFA) increased their likeliness to produce a response to fentanyl (**Fig. 4I**). This indicates that pain enhances the ability of fentanyl to elicit VTA DA neuron activity over time, selectively in males. Initially, the magnitude of fentanyl-evoked responses was attenuated in males with pain relative to males without pain (week 1, **Fig. 4J**). However, this effect was reversed throughout the course of self-administration (**Extended Data Fig. 4F**). Specifically, the maximum dF/F evoked by self-administered fentanyl during week 3 corresponded with pain-dependent effects found on fentanyl intake during week 3 of self-administration (**Fig. 1**). Pain increased peak dF/F in males without affecting peak transients in females (**Fig. 4K**) which parallels pain-dependent increases in fentanyl intake found in males at this time point (**Fig. 1**). Moreover, the average size of the initial response after the lever press was correlated with fentanyl intake during the session in males (**Fig. 4L)** but not females (**Extended Data Fig. 4H)** suggesting that phasic, fentanyl-evoked VTA DA cell responses drive fentanyl reinforcement in a sex-specific manner. Based on the current results and existing evidence indicating the relevance of VTA DA activity in reward processing and motivation^21,45^, peak dF/F triggered by self-administered fentanyl infusions likely encode motivational salience of opioids.

### Protracted increases in fentanyl-evoked VTA DA neuron activity are necessary to increase fentanyl intake in males in the setting of pain

To assess whether the protracted increased responsiveness of VTA DA neurons to fentanyl is required for inflammatory pain to increase fentanyl consumption in male rats, we used a chemogenetic approach in conjunction with fiber photometry and fentanyl self-administration (**Fig. 5A and Extended Data Fig 5-6).** Male TH-Cre+ rats were bilaterally injected with a Cre-dependent inhibitory DREADD (AAV5-hSyn-DIO-hM4D(Gi)-mCherry) or control virus (AAV5-hSyn-DIO-mCherry) in the VTA. In addition, rats were unilaterally injected with the same Cre-dependent calcium sensor as before (pGP-AAV-syn-FLEX-jGCaMP7c-WPRE) and implanted with an optic fiber in the VTA. All rats received hind-paw injections of CFA and IV catheters prior to undergoing fentanyl self-administration following a timeline identical to the previous experiments except that CNO (1 mg/kg, i.p.) or saline (SAL; 1 mL/kg, i.p.) was administered 20 min prior to the start of each session during week 3 (**Fig. 5A**). Tonic and phasic VTA DA activity was measured throughout self-administration as described above.

**FIGURE 5:**
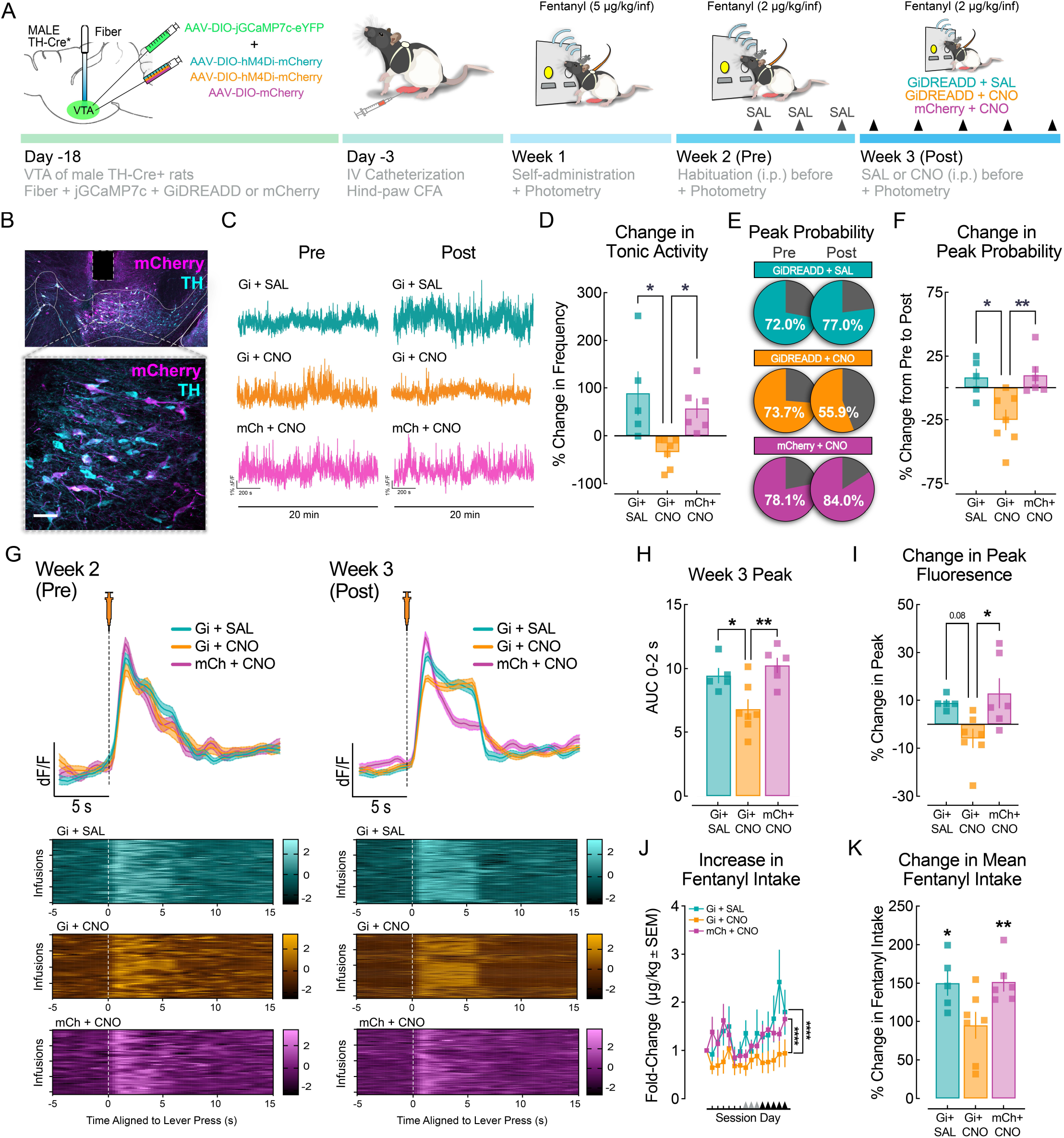
Protracted increases in VTA DA cell calcium dynamics are necessary for pain-dependent increases in fentanyl consumption in males. **A,** Schematic depicting experimental timeline. Male TH-Cre+ rats were injected in the VTA with cre-dependent jGCaMP and inhibitory DREADDS (Gi) or control virus (mCh). Two weeks later, rats received hind-paw injections of CFA before fentanyl self-administration. The last three days of week 2, rats were habituated to the injection procedure with injections of saline (SAL, i.p.). During week 3, rats expressing Gi-DREADDs received either SAL (GI-SAL) or CNO (Gi-CNO) and rats expressing mCh received CNO (mCh-CNO) before the self-administration session. **B,** Representative images of Gi-DREADD (mCherry) expression colocalized with tyrosine hydroxylase (TH) and fiber placement in the VTA of male TH-Cre+ rats. **C,** Representative traces of tonic calcium transient events throughout the 2-hr self-administration session. **D,** Percent change in tonic activity (frequency) after treatment with CNO (1 mg/kg, i.p.) or SAL (1 mL/kg, i.p.) (one-way ANOVA, F_(2,15)_=6.495, P=0.0093, Tukey’s post-hoc test, *P<0.05). **E,** Probability of fentanyl-evoked peaks during week 2 (Pre) or during week 3 (Post) in rats expressing Gi-DREADDs treated with SAL (blue; n=5 rats, 82-103 infusions) or CNO (orange; n=7 rats, 56-161 infusions) or rats expressions mCherry and treated with CNO (magenta; n=6 rats, 55-109 infusions). **F,** Percent change in success rate from week 2 to week 3 (one-way ANOVA, F_(2,15)_=7.878, P=0.0046, Tukey’s post-hoc test, *P<0.05, **P<0.01). **G,** Mean ΔF/F signal aligned to lever press (syringe) during week 2 (Pre, *left*) and week 3 (Post, *right*) of fentanyl self-administration (n=5-7 rats; data expressed as mean ± SEM) and heatmaps for each infusion obtained below. **H,** Area under the curve (AUC) 0-2 s after fentanyl lever press is lower in Gi-CNO rats compared to Gi-SAL or mCh-CNO (One-way ANOVA, F_(2,15)_=8.360, P=0.0036, Tukey’s post-hoc test, *P<0.05, **P<0.01). **I,** Percent change in peak fluorescence after treatment (One-way ANOVA, (F_(2,15)_=5.439, P=0.0167, Tukey’s Post-hoc test, *P<0.05). **J,** Ratio of fentanyl intake normalized to body weight relative to first day of self-administration is lower in Gi-CNO rats compared to Gi-SAL or mCh-CNO (RM one-way ANOVA, F_(2,28)_=22.10, P<0.0001, Holm-Sidak’s multiple comparisons test, ****P<0.0001). **K,** Percent change in fentanyl intake from week 2 to 3 does not increase in Gi-CNO rats (one-sample t-test, *P<0.05, **P<0.01).

Chemogenetic inhibition of VTA DA neurons during the third week of fentanyl self-administration reduced tonic VTA DA cell activity (**Fig. 5B-D**). In the previous experiment, we found that phasic calcium transients from VTA DA cells evoked by self-administered fentanyl were enhanced during week 3 in males with pain, an effect that was correlated with increased fentanyl intake (**Fig. 4**). Chemogenetic inhibition of VTA DA cells during week 3 prevented this time-dependent enhancement in VTA DA cell responsiveness as indicated by a reduction in the probability of fentanyl to elicit responses between weeks 2 and 3 (**Fig. 5E-F**). Moreover, this manipulation prevented the development of high-amplitude, phasic VTA DA cell calcium transients which was revealed by a lack of increase in fentanyl-evoked peak fluorescence between weeks 2 and 3 (**Fig. 5G-I**). Chemogenetic inhibition of VTA DA neurons also prevented increases in fentanyl intake in males with pain (**Fig. 5J**) which was significantly less than control groups during week 3 (**Fig. 5K**). These findings indicate that the growth in phasic VTA DA responses to fentanyl in males with pain is necessary for increased fentanyl consumption.

## DISCUSSION

Pain alters function of the mesolimbic DA system which can lead to altered reward processing and motivational states^21,23,46,47^ due to deficits in VTA DA neuron excitability^20,48,49^ and consequent DA release in the NAc^16,17,50^. As such, opioid reward - encoded by this mechanism - may be perceived differently under conditions of pain. Historically, misconceptions that pain had sparing effects on opioid misuse liability^51^ triggered initial waves in the ongoing opioid crisis^52^ – which is currently exacerbated by the availability of fentanyl, a synthetic opioid up to 100x more potent than morphine^53^. An overlooked but persistent factor in the opioid crisis is the disproportionate number of males affected ^4,29,30,54–57^. Gender-specific tendencies toward negative outcomes associated with long-term prescription opioid use^6^ raise the need to elucidate sex differences in neural mechanisms underlying pain effects on opioid use.

Here, we establish a role for VTA DA neuron activity in mediating sex-specific effects of pain underlying maladaptive patterns of opioid use. We identified sex-specific and time-dependent effects of inflammatory pain on opioid reward seeking and corresponding activation of VTA DA neurons. In males, pain reduced tonic activity of VTA DA neurons indicated by a reduction in baseline frequency, consistent with previous reports from our lab^23^. However, phasic, fentanyl-evoked VTA DA calcium transients were bidirectionally modulated by pain in males. During the first week of self-administration pain reduced dF/F peak in males, which normalized during week 2, before increasing during week 3. Importantly, the trajectory of growth in time-locked fentanyl-evoked responses of VTA DA neurons aligned with rates of fentanyl use such that males in pain had higher intake during week 3 compared to females or males without pain. Recent research found similar effects of pain on tonic and phasic dopamine release in the NAc using fast scan cyclic voltammetry in male rats under light anesthesia ^58^. In this study, cornea application of capsaicin reduced tonic DA release but enhanced electrically evoked phasic DA release, and this effect diminished as the pain resolved^58^. The inverse relationship between tonic and phasic DA release may be mediated by reduced feedback inhibition via DA D2 receptors (D2R) which function as autoreceptors^59–64^. This is supported by a wealth of evidence demonstrating pain-induced deficits in D2R function and expression in the striatum, which includes the NAc^17,65–72^. As such, pain-dependent reductions in D2R binding may permit higher levels of phasic DA release, but it is unclear if this mechanism can explain the lack of pain effects on VTA DA activity in females. Some studies report a lack of sex differences in striatal D2R binding^73–75^, but this has not been evaluated in the context of pain. Despite similarities in D2R binding, clinical reports have found that the magnitude of amphetamine-evoked DA release is higher in men^73^ which contrasts with preclinical reports of higher cocaine-evoked DA in females^76^. Thus it will be important for future studies to characterize the precise cellular mechanisms by which sex differences in phasic DA responses to opioids emerge.

Here, we found that the ability of pain to enhance opioid intake was dependent on pain-facilitated opioid-evoked responses in VTA DA neurons. Pain disrupts function of mesolimbic DA pathways which can lead to clear deficits in motivation for natural rewards ^22,23,77^. However, the effects of pain on opioid motivation are more complex. Our findings here indicate that pain increases fentanyl intake, but only in males after two weeks of training. Consistent with this, previous reports indicate that the ability of pain to affect opioid intake corresponds to the chronicity of pain, dosing, and the duration of exposure. Our findings align with others that demonstrate fentanyl intake may follow the time-course of pain progression, peaking 2-3 weeks after pain onset and declining once pain has subsided^78–80^. In some instances, pain can reduce opioid self-administration at earlier time points^81^. It is possible that we failed to detect initial reductions in opioid intake due to the higher dose of fentanyl (5 μg/kg/infusion) used during the first week of training. This would align with our past research demonstrating inflammatory pain reduces heroin intake at low doses and increases intake at high doses^20^. In this study, we also found that VTA DA neurons were less excitable shortly after the induction of pain due to increased inhibitory drive^20^. Similarly, we found that fentanyl-evoked VTA DA neuron activity was attenuated in males during the first week of self-administration and expand on these by demonstrating a time-dependent increases in fentanyl-evoked calcium dynamics. Recent findings from our lab identified the rostromedial tegmental nucleus (RMTg) as the critical source of pain-mediated VTA DA neuron inhibition, a circuit necessary to produce motivational deficits in natural reward seeking^23^. The RMTg is also the most opioid-sensitive source of inhibition in the VTA^12,82^ and undergoes plasticity in a sex-specific manner^83^. As such, the RMTg→VTA pathway may represent the means by which pain dynamically influences VTA DA neuron activity underlying sex-specific patterns of maladaptive opioid use. Future studies delineating the sex-specific role of this pathway in pain-dependent effects on opioid use will further our understanding of the circuit specific mechanisms underlying the current findings.

The effects of pain on fentanyl-evoked VTA DA neuron activity are likely relayed at downstream targets important for reward and encoding motivational salience, like the NAc ^84–87^. Our past research showed that inflammatory pain bidirectionally modulates DA transmission in the NAc, dose-dependently, after passive heroin exposure^20^. These effects were correlated with similar effects on the motivation for heroin self-administration, increasing for higher and decreasing for lower doses^20^. The present results add to these findings by detecting the effects of pain on fentanyl self-administration and real-time associated VTA DA neuron activity. This is critical from a translational perspective given that other studies examining sex differences in opioid self-administration show inconsistent results. In this regard, most studies report higher rates of opioid self-administration in females compared to males^88–100^ while others fail to capture any sex differences^99–102^. Notably, one study found that males self-administer more oxycodone when demand is low (FR1) and this modest effect was on the number of infusions obtain rather than lever responses, suggesting that males were only slightly more efficient during the 6-s timeout periods^103^. This existing bias in preclinical studies of opioid use is - in part - attributed to historically sex/gender biased research, but points to a critical gap in basic research.

Population-based and qualitative evidence indicates higher rates of opioid misuse liability in men based on negative outcomes associated with opioid use, particularly, overdose deaths^6,31^. The inability of preclinical research to recapitulate similar sex differences is unlikely due to experimental parameters but rather, interplay with a tertiary variable. Most opioid users report ‘pain’ as their primary reason for opioid misuse^104^. Despite documented sex differences in pain prevalence^24,105^, nociception ^106,107^, opioid sensitivity/analgesia^26,28^, and tolerance^108^, - no preclinical research to date has evaluated chronic pain effects opioid use in a sex specific manner. Our results are novel in that they begin to address a major gap in our current understanding of the relationship between pain and opioid misuse. To advance pain treatment in the face of the opioid crisis, it will be critical for opioid and pain research to detail sex-specific mechanisms underlying patterns of opioid abuse in the context of pain.

## Acknowledgements

We thank Dr. Nikolas Massaly, Dr. Tamara Markovic, and Kristine Yoon for their theoretical contributions to the conceptualization of this project and assistance with training. We thank Justin Meyer for breeding the colonies used in these experiments and for general support throughout the experiments. Finally, we thank Jordan Nagai and Hasan Maqbool for their technical assistance. This work was supported by US National Institutes of Health (NIH) grants DA726129 (JH), DA041781 (JM), DA042581 (JM), DA042499 (JM), DA041883 (JM), and DA045463 (JM), W.M. Keck Fellowship (JH), and NARSAD Independent Investigator Award from the Brain and Behavior Research Foundation (JM).

## Author Contributions

Conceptualization: JH and JM; Methodology: JH, JA, RT, and JM; Formal analysis of all data: JH, JA, RT, and JM; Photometry acquisition and data extraction: JH and JA; Operant self-administration; JH and JA; Von Frey: JH and JA; Immunohistochemistry, microscopy, and cFos counts: JH, JA, and RT; Surgeries: JH and JA; Writing of original draft: JH and JM; Writing (review and editing): JH, JA, RT, and JM; Funding acquisition: JH and JM; Resources: JM; Supervision: JH and JM.

## METHODS

All procedures were approved by Washington University and the National Institute on Drug Abuse (NIDA) Animal Care and Use Committee in accordance with the National Institutes of Health Guidelines for the Care and Use of Laboratory Animals.

### Animals

Adult male and female Long Evans wild-type, TH-Cre (250–350 g) were used for this study. All animals for behavioral experiments were 8–10 weeks old at the beginning of the experiments. Rats were single-housed on a 12/12-h dark/light cycle (lights on at 7:00) and acclimated to the animal facility holding rooms for at least 7 days before any manipulation. The temperature for the holding rooms of all animals ranged from 21 to 24 °C while the humidity was between 30 and 70%.

All experiments were performed during the light cycle. Rats received food and water ad libitum until 2 days before starting the behavioral studies, when mild food restriction (25 or 30 g of rat chow per day for females or males, respectively) was implemented to maintain weights within 50 g of starting weights and continued until the end of the experiments.

### Surgeries

All surgeries were performed under isoflurane (2.5/3 minimum alveolar concentration) anesthesia using appropriate sterile aseptic techniques.

### Intracerebral injections

For chemogenetic inhibition of VTA DA neurons and fiber photometry recordings of VTA DA neuron activity, TH-Cre rats were bilaterally injected with AAV5-hSyn-DIO-hM4D(Gi)-mCherry, AAV5-hSyn-DIO-mCherry and/or pGP-AAV-syn-FLEX-jGCaMP7c variant 1513-WPRE^109^ (all viruses injected at 1.5–2.5 × 10^12^ transducing units per ml (0.5 μl per side), Addgene), and implanted with an optic fiber (Doric Lenses) in the VTA (stereotaxic coordinates for viral injections from Bregma: anterior–posterior (AP) = −5.28 mm, medial–lateral (ML) = ±3.0 mm, dorsal–ventral (DV) = −8.45 mm at a 15° angle; stereotaxic coordinates for optic fiber from Bregma: AP = −5.28 mm, ML = ±0.78 mm, DV = −7.85 mm from the skull surface, World Precision Instruments). The optic fiber implant was secured to the skull using three sterile bone screws and a dental cement head-cap (Lang Dental). To avoid post-surgical complications and to minimize pain, animals received injections of enrofloxacin (8 mg/kg, s.c.) and carprofen (5 mg/kg, s.c.) before surgery and a chewable bacon-flavored rimadyl tablet (2 mg/tablet; Bio-Serv) 0, 24, and 48 h after surgery. Rats were given a 2-3 week recovery period before undergoing jugular catheterization.

### Jugular Catheterization

To permit intravenous fentanyl self-administration, jugular catheters constructed in house were surgically implanted into the right jugular vein as previously described^110,111^. Silastic tubing exiting between the scapulae was connected to a single-channel silicone vascular access harness (Instech). As described above, enrofloxacin (8 mg/kg, s.c.) and carprofen (5 mg/kg, s.c.) and a chewable bacon-flavored rimadyl tablet (2 mg/tablet; Bio-Serv) were administered 0, 24, and 48 h after surgery. Catheter patency was maintained with daily flushing of 0.3 mL sterile saline containing gentamicin (1.33 mg/mL, i.v.).

### CFA injections

At the end of the jugular catheterization procedure, rats were maintained under anesthesia and intraplantar injections of sterile saline or CFA (Thermo Fisher) were administered to the right hindpaw at volumes of 120 or 150 μL for females or males, respectively. The general behavior (feeding, drinking and locomotion) of the animals was monitored throughout the duration of experiments.

### Fentanyl Self-Administration

To mitigate potential confounds in the acquisition of self-administration^112^, rats did not undergo any instrumental training prior to the start of fentanyl self-administration (SA). Fentanyl SA training was initiated 72 hours after catheterization and hind paw injections with CFA or saline in operant-conditioning chambers (Med Associates, MED, PC 5 software) equipped with two retractable levers positioned on the right-hand wall with two cue lights positioned above each lever. Each week of self-administration consisted of 5 consecutive training days (2 days off) under a fixed-ratio 1 (FR1) schedule of reinforcement. During the 2-h self-administration sessions, both levers (active and inactive) were extended and responses on the active lever resulted in illumination of the cue light for 5 s and simultaneous delivery of a fentanyl infusion (100 μL, i.v.) at a dose of 5 or 2 μg/kg/infusion during training weeks 1 or 2/3, respectively. These doses were determined in preliminary experiments based the minimally analgesic dose (2 μg/kg) and maximally effective dose devoid of any effects on locomotor activity. Presses on the inactive lever were recorded but had no programmed consequences. Fentanyl SA training lasted three weeks (15 sessions) independent of performance to maintain analogous CFA/SAL inoculation periods across groups. Upon completing three weeks of fentanyl SA, rats continued training the following week on (a) progressive ratio (PR) and fentanyl dose-response sessions or (b) extinction and cue-induced reinstatement.

### Progressive Ratio and Dose-Response Tests

For PR and dose-response sessions, rats underwent 2 daily 2-h fentanyl SA sessions (2 μg/kg/infusion) under a PR schedule of reinforcement during which each correct response requirement increased exponentially during the subsequent trial (5 × *e*^(0.2 × infusion number)^ – 5 rounded to the nearest integer resulting in the following PR steps: 1, 2, 6, 9, 12, 15, 20, 25, 32, 40, 50, 62, 77, 95….). The next day, rats continued with 2 daily FR1 sessions (as described above) to mitigate potential confounding effects on subsequent dose–response sessions. Next, rats underwent 2 consecutive within-session dose–response tests. One of three doses of fentanyl (2, 3, 5 μg/kg/infusion) was available on a FR1 reinforcement schedule for 1 h during the session, with a 5 min resting period between doses. Doses were altered by varying the infusion duration. The presentation of the doses was made in an ascending or descending manner and the order of dose presentation between the two days was counterbalanced; thus, half of the rats followed ascending order (2, 3, 5 μg/kg/infusion) during the first dose-response test and descending order (5, 3, 2 μg/kg/infusion) during the second dose-response test, while the other half of rats were exposed to descending then ascending orders during the sessions.

### Extinction and Reinstatement

For rats that underwent extinction and reinstatement, extinction training occurred during 7 daily one-hour sessions in which fentanyl-paired cues and fentanyl infusions were withheld. Both levers were extended during the extinction sessions and responses were recorded but had no programmed consequences. After 7 extinction training sessions, rats underwent a 2-hour reinstatement test in which correct lever responses resulted in illumination of the cue light over the correct lever (as in self-administration) except that fentanyl reinforcement was withheld.

### Tests for Nociception, Antinociception, and Inflammation

To assess baseline nociception (i.e. mechanical hyperalgesia) induced by CFA injections, paw withdrawal thresholds (PWT) were obtained using an electronic Von Frey Anesthesiometer (IITC Life Science). Animals were placed in plexiglass chambers on top of a galvanized steel mesh shelf to permit access to the rats’ paws from underneath. The anesthesiometer was used to provoke a flexion reflex followed by a flinch response, and the mechanical threshold pressure in grams (PWT) was recorded. Rats were habituated to the test chambers and Von Frey procedure for at least 1 h at least one day prior to undergoing self-administration. On assessment days (every other day of self-administration), rats were placed in the plexiglass chambers at least 2 h after completing the self-administration session to mitigate potential lasting analgesic effects of fentanyl. Rats were acclimated to chambers for 30 min prior to undergoing the procedure. Once acclimated, baseline measurements of mechanical sensitivity were obtained in triplicates for each paw at 5-min intervals, alternating between the injected (right) and non-injected (left) paw. Following baseline measurements, antinociceptive responses were derived from a single non-contingent injection of fentanyl (2 μg/kg, i.v.) that was delivered through the indwelling catheter. Mechanical sensitivity was assessed in the same manner except that, measurements were collected in duplicates from each paw within 5 min of the injection to accommodate the short half-life of fentanyl. Finally, the rats were removed from the chambers and a caliper was used to assess the maximal dorsal-ventral hind paw thickness (in mm) of the left and right paws before animals were returned to the home cage.

### Immunohistochemistry

Following each behavioral experiment, rats were transcardially perfused with PBS followed by 4% paraformaldehyde. For the detection of cFos and to verify viral expression, rats were perfused 2 hr after injection of either CNO (1 mg/kg, i.p.) or saline (1 mL/kg, i.p.) to permit maximal neuronal activation. Brains were collected and kept at 4 °C for 24 h in 4% paraformaldehyde for post-fixation, followed by at least 72 h incubation in 30% sucrose solution. Isopentane was used to flash-freeze brains, which were cut in 40-μm coronal slices using a cryostat (Leica CM 1950). Free-floating sections containing the VTA were washed in PBS and blocked with normal donkey serum (Millipore, S30) and 0.3% Triton–0.01 M PBS (PBST) and incubated overnight in 3% normal donkey serum and 0.3% PBST containing primary antibodies for GFP (1:2000, chicken anti-GFP, ab13970, Abcam), and/or TH (1:2000; mouse anti-TH, MAB318, Millipore), and/or mCherry (1:500; rabbit anti-mCherry, ab167453, Abcam or 1:1000; mouse anti-mCherry, ab1C51, Novus Biologicals), and/or cFos (1:1000; rabbit anti-cFos, ab214672, Abcam). The next day, sections were washed with PBS and incubated (2 h) with the appropriate secondary antibodies diluted in PBS (donkey anti-chicken A488 or donkey anti-rabbit A488; donkey anti-mouse Cy3 or donkey anti-rabbit Cy3; donkey anti-mouse DyLight405). All secondary antibodies were diluted 1:250 and were from Jackson ImmunoResearch.

A Leica DMR microscope was used to process images (×5, ×10 and/or ×20 magnification) and detect viral expression. ImageJ 1.53a software was used to manually quantify the number of labeled cells for combined chemogenetics experiments and cFos activation. For viral expression verification, z-stacks spanning the VTA were collected using a 10X objective from −4.80 AP to −6.12 AP. For cFos quantification, z-stacks of the VTA were collected using a 20x objective and 3 VTA slices of equivalent distance from Bregma between −4.80 AP and −6.12 AP were quantified per animal. Cell counts were done by two independent researchers that were blinded to the experimental results and treatments.

### In Vivo Wireless Photometry Acquisition

For fiber photometry experiments, rats underwent the same behavioral protocol described for fentanyl self-administration. Fiber photometry recordings were made throughout the entirety of the 2-h self-administration training sessions on at least one day per week to capture various timepoints throughout self-administration. To overcome interference with the infusion line and patch cable, we used a wireless photometry device (TeleFipho, Amuza Inc.). An LED generated blue light was bandpass-filtered (445-490 nm) to excite GCaMP7c and emission fluorescence was detected by an internal photodiode detector detecting green light (bandpass filtered at 500-550 nm). An internal DC amplifier transmitted the data (16-bit arbitrary unit; AU) to a wireless receiver (TeleFipho, Amuza Inc.) and the data was extracted in real-time using TeleFipho software (Amuza Inc.) at a sampling rate of 100 Hz. Prior to the first day of recording, the wireless headstage was secured to the optic fiber and the offset was set to 90° and LED power was adjusted within the range of optimal output signal (30-40 k, AU), but not to exceed 36 μW. The settings were recorded and maintained for each individual animal across each recording days. Fiber photometry signals were time-locked with correct lever responses through an analog output from MedAssociates transmitted through a BNC cable to the wireless receiver and voltage signals were extracted in real time using a second channel in TeleFipho software at a sampling rate of 100 Hz.

The first 5 minutes of each recording session were removed the analyses to mitigate effects of baseline drift in signal due to slow photobleaching artifacts. The reliability of fluorescent signals over time were verified using a simple linear regressions and two-tailed Pearson’s correlations using the raw mean fluorescent output values (arbitrary units, a.u.) at the session start and/or end, percentage of signal decay due to photobleaching within each session (session end output / session start output x 100) and LED power (μW) associated with each recording session. Residual effects of baseline drift were corrected for in session frequency analysis by fitting a double-exponential curve to the raw trace and subtracted. Baseline correction of session data analyzed for time-locked events were fit to a single exponential curve, subtracted, and trial-trace data was extracted and sampled at 2 Hz using the pMAT software suite in Matlab. After baseline corrections, photometry traces were normalized to units of *ΔF/F* using the median fluorescence of the entire session or trial for subtraction and division. Custom Matlab scripts were developed for analyzing tonic calcium transient event rates (Hz) for individual sessions. Events throughout the 2-h session were determined by identifying local maxima in the Δ*F/F* photometry signal that resembled the waveform and temporal dynamics of GCaMP calcium transients. The built-in Matlab function findpeaks was used to identify local maxima that were at least 1.2x Δ*F/F*% in prominence and at least 150 ms in width at half prominence (**Fig. 4B**). For quantification of fluorescence during lever pressing, the Matlab function findpeaks was used to identify events at −1 to +6 s relative to each lever press (Time=0) with at least 1.2x Δ*F/F*% in prominence and at least 500 ms in width at half prominence. Peak fluorescence in response to fentanyl was calculated at the maximum *ΔF/F* at −1 to +6 s relative to reward delivery in trials were events were detected. The response rate (%) was determined as the number of trials (fentanyl-reinforced lever presses) with detectable events divided by the total number of trials times 100. Area under the curve for detected events was calculated in 2-s bins from −4 to +8 s using the trapezoidal rule. Alpha was set to 0.05 for all analyses.

### Experimental design and statistics

All the experiments were successfully replicated at least twice, including each treatment condition, to prevent a nonspecific day/condition effect. Treatment groups were randomly assigned (CFA or Saline) to animals before the start of the first day of self-administration or counter-balanced (CNO or Saline) based on self-administration history for DREADD manipulations. After assessing the normality of sample data using D’Agostino and Pearson tests and Shapiro–Wilk tests, statistical significance was taken as \**P* < 0.05, \*\**P* < 0.01, \*\*\**P* < 0.001 and \*\*\*\**P* < 0.0001, as determined by one-way ANOVA, repeated-measures (RM) 2-way ANOVA, 2-way ANOVA, or 3-way ANOVA followed by Tukey’s or two-tailed Sidak’s post hoc test, Friedman’s test, two-tailed unpaired *t*-test, two-tailed paired *t*-test, one-sample *t-*test, Kruskal–Wallis test, two-tailed Dunn’s multiple comparisons test, two-tailed Wilcoxon test, two-tailed uncorrected Dunn’s test, Dunnett’s multiple comparisons test, Kolmogorov–Smirnov test, or two-tailed Mann–Whitney for unpaired values as appropriate. No outliers were excluded from any of the studies presented in this manuscript. Statistical analyses were performed using GraphPad Prism 8.1.0 and Matlab. Data collection and analysis were performed blinded to the conditions of the experiments.

