## Extended Data Figures for "Time-dependent enhancement in ventral tegmental area dopamine neuron activity drives pain-facilitated fentanyl intake in males"

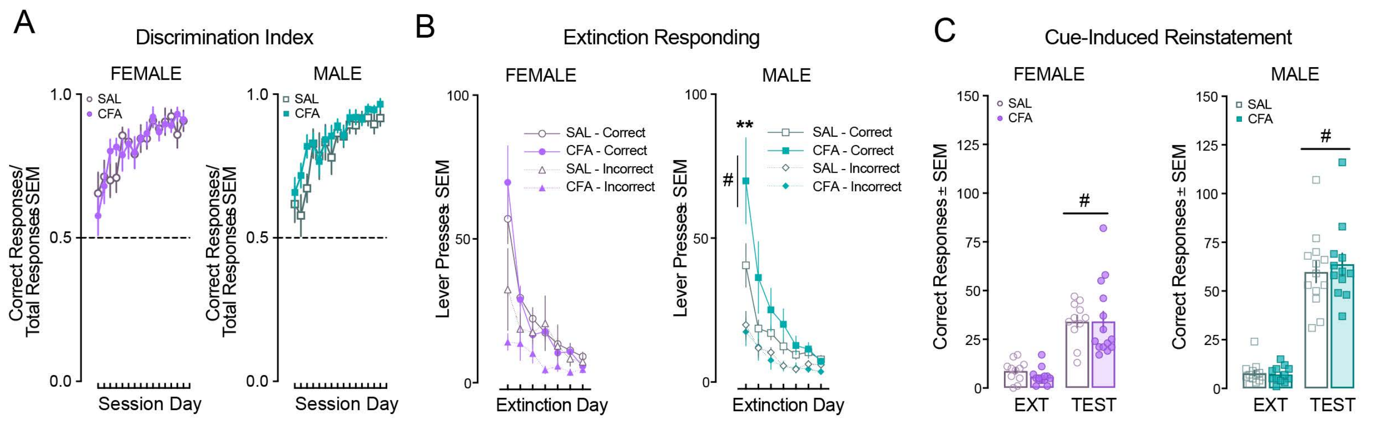


**Extended Data Figure 1: A,** Female (*left)* and male *(right)* discriminate between levers after the first day of self administration as indicated by discrimination indices >0.5 (correct responses/total responses) (2way ANOVA females, time: $F_{(14,448)}$=10.46, P<0.0001; 2way ANOVA males, time: $F_{(14,420)}$=12.92, P<0.0001). **B,** Females (*left)* respond similarly during extinction training (Correct RM 2way ANOVA, time: $F_{(6,144)}$=35.06, P<0.0001; Incorrect, time: $F_{(6,126)}$=3.941, P=0.001. Males (*right)* with pain increase responding on the fentanyl-paired lever on the first day of extinction (Correct RM 2way ANOVA, time x treatment: $F_{(6,138)}$=2.237, P=0.043; Sidak's Post hoc, **P=0.007; incorrect, time: $F_{(6,138)}$=11.63, P<0.0001). **C,** Pain does not effect cue-induced reinstatement in females (*left)* or males (*right)*(female RM 2way ANOVA, cue: $F_{(1,23)}$=61.63, #P<0.0001; male RM 2way ANOVA, cue: $F_{(1,23)}$= 83.2, #P<0.0001).

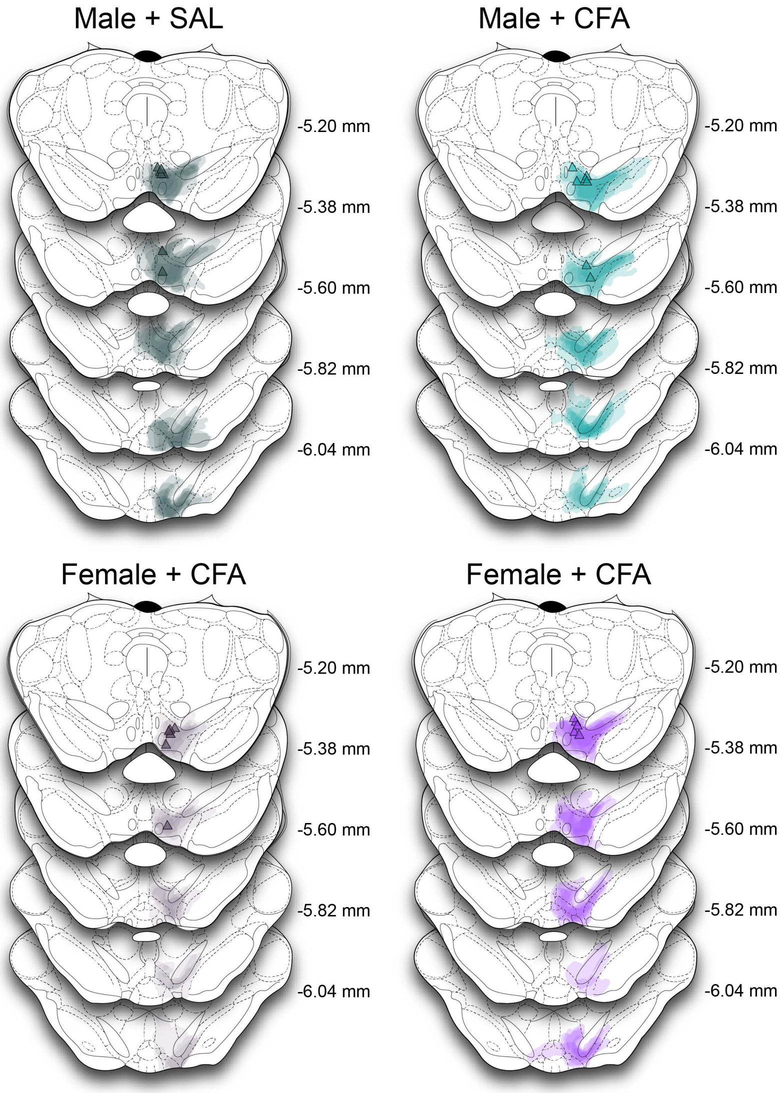


**Extended Data Figure 2**: GCaMP viral expression for each group with the most ventral point of the fiber indicated in triangles. Coordinates relative to bregma.

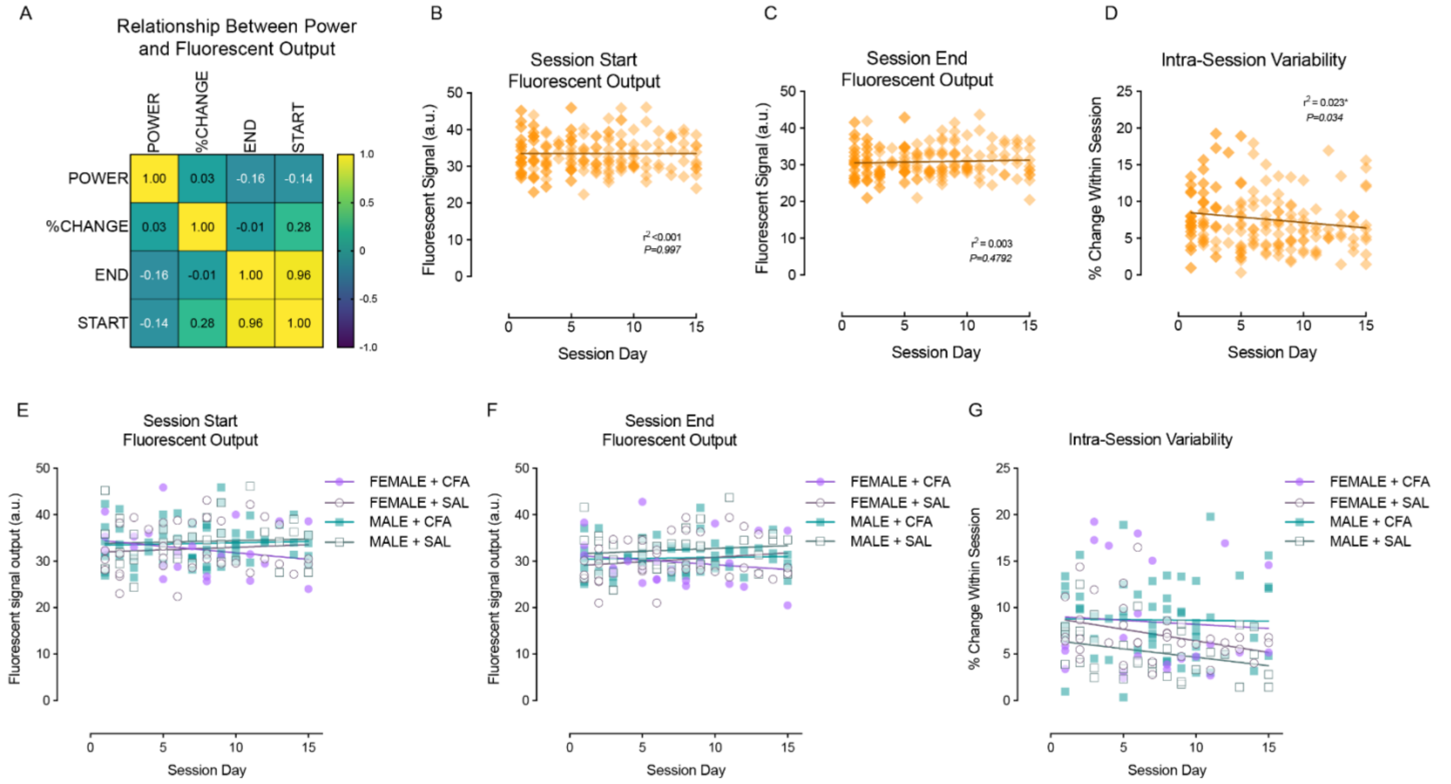


**Extended Data Figure 3: GCaMP fluorescence is stable over time. A,** LED power is not correlated with differences in the magnitude of GCaMP fluorescent within-session photobleaching (%CHANGE), ending output signals (END) or starting output signals (START) (Pearson's correlation coefficient plotted on heatmap). **B,** GCaMP fluorescence at the start of the session measured in arbitrary units (a.u.) in Telefifo software is stable over 15 self-administration sessions (i.e. 3 weeks) (Simple linear regression, $F_{(1,191)}<0.0001$, P=0.997; slope=-0.0004, $r^2<0.0001$). **C,** GCaMP output at the end of the 2-hr recording session is stable up to three weeks (Simple linear regression, $F_{(1,191)}=0.514$, P=0.479; slope=0.054, $r^2=0.003$). **D,** Intra-Session variability determined as the percent change in signal decay within the 2-hr session ([start output-end ouput]/start output*100) was slightly reduced over time (Simple linear regression: $F_{(1,191)}=4.58$, *P=0.034; slope=-0.149, $r^2=0.023$). **E,** There were no differences between groups in the starting output over time (Simple linear regression, $F_{(3,144)}=0.6007$, P=0.6156). **F,** Treatment groups exhibited similar and stable output at the end of the session over time (Simple linear regression, $F_{(3,144)}=0.7685$, P=0.5141). **G,** Intra-session variability in signal decay caused by photobleaching was also similar between groups over time (Simple linear regression, $F_{(3,144)}=0.5229$, P=0.6672).

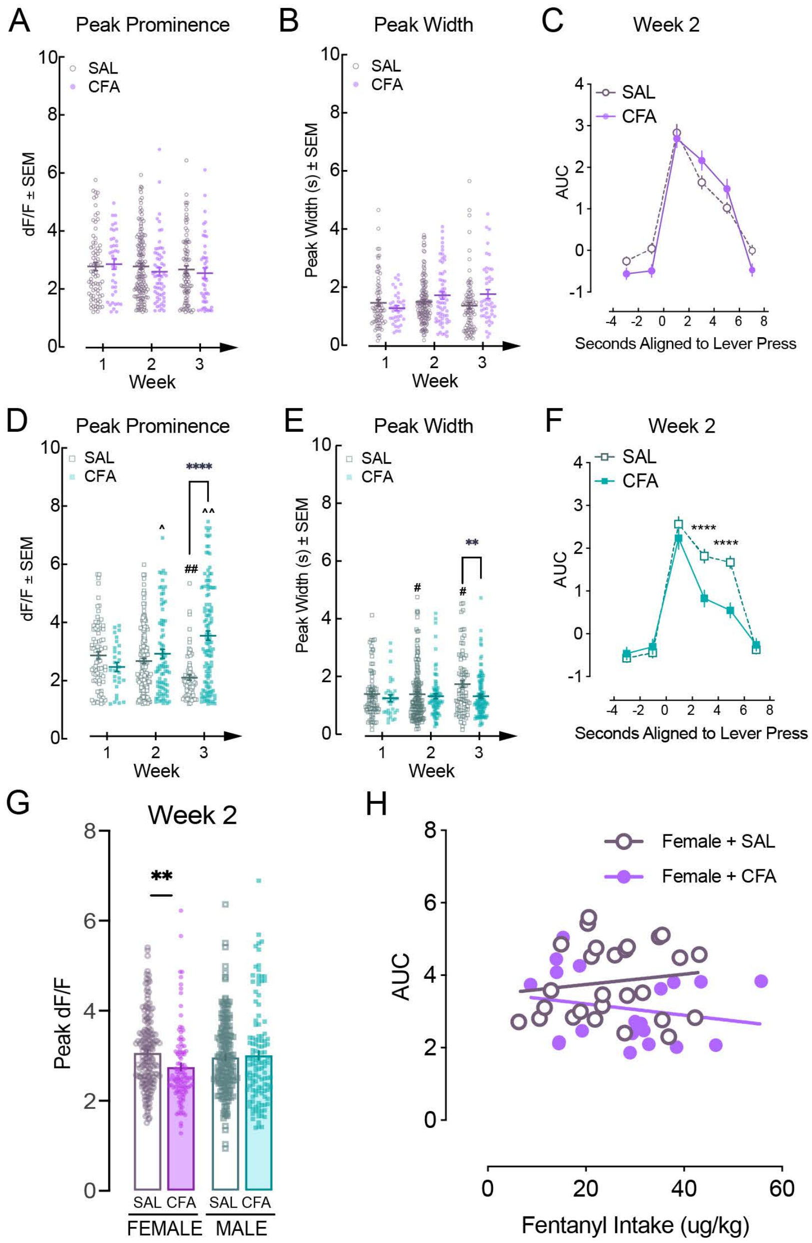


**Extended Figure 4: Time-dependent effects on fentanyl-evoked phasic VTA DA neuron calcium transients. A,** In females, peak prominence (rise above baseline) is similar between female groups across the three weeks of self-administration (2way ANOVA, ns). **B,** In females, peak width (duration) is similar between groups across the three weeks of self-administration (2way ANOVA, ns). **C,** Area under the curve (AUC) measured in 2-s intervals relative to lever presses were similar over time in females during week 2 (RM 2way ANOVA, ns). **D,** Peak prominence of males without pain decreases each week of fentanyl self-administration (2way ANOVA, time x treatment: $F_{(2,535)}$=13.79, P<0.0001; treatment: $F_{(1,535)}$=18.75, P<0.0001; Sidak's Post hoc, ##P<0.01 relative to week 1) while peak prominence in males with pain increases over time (Sidak's Post hoc, ^P=0.0219, ^^P=0.0032) and peak prominence is higher in CFA males than SAL males (Sidak's Post hoc, ****P<0.0001). **E,** Peak width is attenuated in males with CFA relative to SAL during week 3 (2way ANOVA, time x treatment: $F_{(2,535)}$=2.112, P=0.1220; time: $F_{(2,535)}$=2.632, P=0.0739; treatment: $F_{(1,535)}$=6.393, P=0.0117; Sidak's post-hoc, **P<0.01) and increases over time in males with SAL (Sidak's post hoc, #P<0.05 relative to week 1). **F,** AUC of fentanyl-evoked calcium transients in males in week 2 show that the effects of CFA shift to latent decreases in AUC (2way ANOVA, time x treatment: $F_{(5,1806)}$=6.494, P<0.0001; time: $F_{(5,1806)}$=109.5, P<0.0001; treatment: $F_{(1,1806)}$=14.07, P=0.0002; Sidak's post hoc, ****P<0.0001). **G,** Pain reduces fentanyl-evoked dF/F in females during self-administration week 2 (2way ANOVA, sex x treatment: $F_{(1,574)}$=5.604, P=0.0183; Sidak's Post hoc, **P=0.008). **H**, No correlation between fentanyl intake and AUC (0-2 s) in females regardless of treatment.

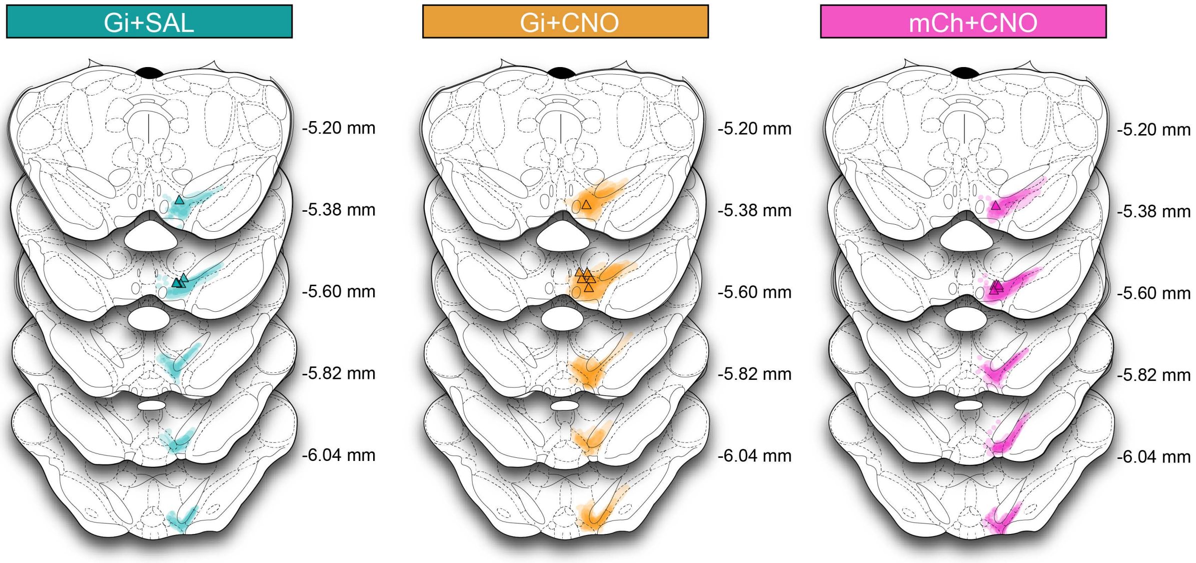


**Extended Data figure 5:** GCaMP viral expression for each group with the most ventral point of the fiber indicated in triangles. Coordinates relative to bregma.

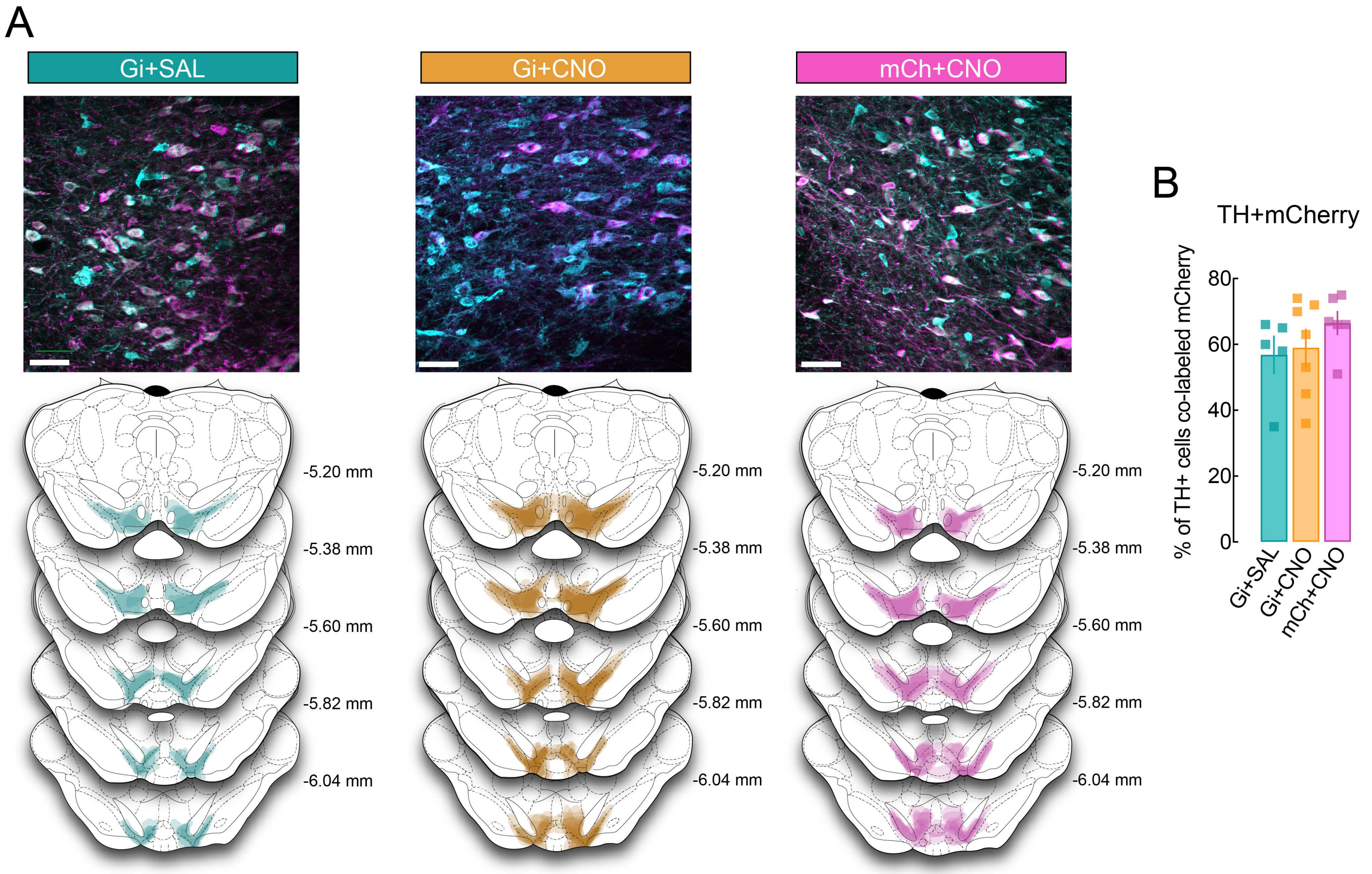


**Extended Data Figure 6: A,** Representative image (top) of viral expression in the VTA (scale bar = 50 µm) of inhibitory DREADDs (Gi) or control virus (mCh) for each group and schematic depicting viral spread. Coordinates relative to bregma. **B,** Percentage of VTA cells expressing tyrosine hydroxylase (TH+) co-localized with mCherry is similar across groups (one-way ANOVA, n.s.).
